# SEQUENCE VS. STRUCTURE: DELVING DEEP INTO DATA-DRIVEN PROTEIN FUNCTION PREDICTION

**DOI:** 10.1101/2023.04.02.534383

**Authors:** Xiaochen Tian, Ziyin Wang, Kevin K. Yang, Jin Su, Hanwen Du, Qiuguo Zheng, Guibing Guo, Min Yang, Fei Yang, Fajie Yuan

## Abstract

Predicting protein function is a longstanding challenge that has significant scientific implications. The success of amino acid sequence-based learning methods depends on the relationship between sequence, structure, and function. However, recent advances in AlphaFold have led to highly accurate protein structure data becoming more readily available, prompting a fundamental question: *given sufficient experimental and predicted structures, should we use structure-based learning methods instead of sequence-based learning methods for predicting protein function, given the intuition that a protein’s structure has a closer relationship to its function than its amino acid sequence?* To answer this question, we explore several key factors that affect function prediction accuracy. Firstly, we learn protein representations using state-of-the-art graph neural networks (GNNs) and compare graph construction(GC) methods at the residue and atomic levels. Secondly, we investigate whether protein structures generated by AlphaFold are as effective as experimental structures for function prediction when protein graphs are used as input. Finally, we compare the accuracy of sequence-only, structure-only, and sequence-structure fusion-based learning methods for predicting protein function. Additionally, we make several observations, provide useful tips, and share code and datasets to encourage further research and enhance reproducibility.

## 1 Introduction

Proteins are the main machinery of *in vivo* biology processes, carrying out a myriad of diverse functions. The relationships between protein sequence, structure, and function underlies cellular biology: *the primary structure (i.e. amino acid (AA) sequence) within a protein determines its 3D structure (fold), which in turn determines its function* (Zhang et al., 2021; Serçinoğlu & Ozbek, 2020). Protein function prediction (PFP) has routinely built on such theory by taking either AA sequences or 3D structures as input and predicted functions as output. As a broad term, protein function refers to both protein annotations (Zhou et al., 2019) with categorical labels and mutation fitness effects (Dallago et al., 2021) with continuous values.

The revolution in high-throughput sequencing (HTS) technologies has resulted in an enormous and ever-increasing number of new protein sequences discovered every year. For example, the number of sequences in UniProtKB^1^ had risen to approximately 190 million by the end of 2020 (uni, 2021). This quantity of high-quality protein sequence data has enabled the tremendous success of deep learning based PFP methods, in particular large-scale protein language models (PLMs) (Rao et al., 2019; Alley et al., 2019; Rives et al., 2021b; Elnaggar et al., 2020; Rao et al., 2021; Yang et al., 2022a). In contrast, determining a protein’s 3D structure via wet-lab experiments is costly and time-consuming: to date, there are fewer than 190,000 structures in the Protein Data Bank (PDB) (Berman et al., 2000). Therefore, it has been impractical to perform PFP by structure-based methods for many proteins due to the lack of experimental structures.

However, recently AlphaFold (Jumper et al., 2021) accurately predicts protein structure from sequence with near-atomic accuracy, enabling an unparalleled expansion in the number of accurate protein structures available. Furthermore, DeepMind and the European Bioinformatics Institute have partnered to create the AlphaFold Protein Structure Database (AlphaFold DB) with 200 million predicted structures (Varadi et al., 2022).

This has naturally raised a question: *are deep learning methods that use AlphaFold-structures as inputs more accurate than AA-sequence-based PLM methods?* We empirically attempt to answer the following sub-questions:

### Q1: How can protein structure data be modeled to better represent various protein properties?

**A:** We explore four graph constructing methods by considering distances at three different granularities: residues, backbone atoms, and backbone and side chain atoms. We also consider a graph that encodes Cb-Cb distances and backbone dihedral angles. We benchmark three representative graph neural network (GNN) architectures (GCN (Kipf & Welling, 2016), GAT (Velickovic et al., 2017), and GIN(Xu et al., 2018)) against these graph constructing methods. We find that the residue-based composition, using the main chain atoms to calculate the distance, direction and orientation of several features, the results on the graph transformer network are better than the above methods.

### Q2: Is the structural data produced by AlphaFold accurate enough to be used for deep learning methods, compared to experimental data?

**A:** We compare the performance of the same GNN model using structure data generated by AlphaFold and from the PDB. Our observations reveal that structures generated by AlphaFold can produce results that are comparable to those obtained from PDB structures. To further improve prediction accuracy, we conduct graph network pre-training using 2 million structures from the AlphaFold database. Our findings indicate that pre-training on these structures does indeed help to enhance the final prediction accuracy.

### Q3: Which learning approach performs the best for PFP tasks, GNN-based structure model or PLM-based sequence model? Is it beneficial to integrate them into one model?

**A:** Our benchmark experiments demonstrate that, despite the intuitive assumption that protein structures have a closer relationship to function, sequence-based PLMs generally outperform structurebased GNN models. However, we also find that combining protein sequence and structure information can improve performance. These observations suggest that standard GNN models may not be the most effective way to encode protein structure data, and that, for now, sequence-based deep learning models continue to dominate the PFP task.

In addition to the above findings, we have conducted additional studies and identified several tricks and factors that can impact protein modeling. We will release our code and data, along with all experimental details, to facilitate implementation. We believe that our observations may significantly surprise the biological community as they challenge the common belief that a protein’s specific shape or structure directly determines its function and that this is more informative than sequence. We also expect that our results will encourage the AI community to design more powerful structure prediction models and more expressive protein representation models, ultimately leading to improved function prediction.

## 2 Related Work

### Protein Function Prediction

Traditionally, protein function determination is carried out either by *in vitro* or *in vivo* experiments, which are limited by time and cost (Clark & Radivojac, 2011). The revolution of HTS has produced a deluge of unmeasured protein data in diverse species and spawned a series of faster and cheaper computational prediction methods (Bileschi et al., 2022; Radivojac et al., 2013; Rives et al., 2021b; Dallago et al., 2021; Pazos & Sternberg, 2004; Hie et al., 2021;Unsal et al., 2022).

BLAST is a classical *in-silico* method that compares protein sequences to sequence databases and find regions of local similarity (Altschul et al., 1990). However, a simple similarity measure with a pre-set threshold is insufficient to assign high-confident protein function. DL-based PFP methods include function prediction from AA-sequence (Rao et al., 2019; Alley et al., 2019; Elnaggar et al., 2020; Dallago et al., 2021; Kulmanov & Hoehndorf, 2020; Meier et al., 2021; Biswas et al., 2021; Gelman et al., 2021; Yang et al., 2022a), 3-dimensional structure (Gligorijević et al., 2021; Smaili et al., 2021; Guo et al., 2022), evolutionary relationships and genomic context (Rao et al., 2021; Engelhardt et al., 2005), and their combinations (Chen et al., 2021; Gligorijević et al., 2021). Here, we mainly restrict our scope to the sequence and structure. Additionally, PFP can be categorized as annotation prediction (AP) (Radivojac et al., 2013; Zhou et al., 2019), which predicts the presence or absence of a functional label, or fitness prediction (FP) (Dallago et al., 2021), which predicts a real-valued fitness measurement.

### Protein Language Model

Researchers have adapted many natural language processing (NLP) methods to model AA sequences, known as protein language models (PLMs) (Bepler & Berger, 2021). Early work UniRep (Alley et al., 2019) trained an autoregressive Recurrent Neural Network (RNN) on 24 million UniRef50 sequences, demonstrating better results than classical bioinformatics methods on PFP. ESM-1b (Rives et al., 2021a) leverages the BERT (Devlin et al., 2018) reconstruction task to represent proteins, and scales the model to 650 million parameters trained on UniRef50. TAPE (Rao et al., 2019) set up a PLM benchmark with 5 downstream tasks, including both sequence-structure and sequence-function predictions and evaluated a BERT-style model trained on Pfam on these tasks.

ProtBert (Elnaggar et al., 2020) further applies several latest language models for building protein representations, including BERT-based, Albert-based, Electra-based, Transformer-XL-based, XLNet-based and T5-based PLMs. After that, MSA Transformer (Rao et al., 2021) extends the BERT task to multiple sequence alignments (MSAs) in order to learn evolutionary information from homologous sequences. It greatly improves performance for structure prediction but has not been tested for function prediction.

### Protein Graph Model

In addition to the above PLM methods, there is some work (Borgwardt et al., 2005; Fout et al., 2017; Yang et al., 2020; Yuan et al., 2022; Miyazaki et al., 2020; Son & Kim, 2021) attempting to directly encode protein structures into representations and use them to perform function predictions; we refer to these as protein graph models (PGMs). GNNs are a natural choice for modeling protein structures. However, PGMs have received relatively little attention due to the unavailability of high-accurate 3D structure data. DeepFRI (Gligorijević et al., 2021) uses a GCN to predict function from backbone structures. (Fout et al., 2017) developed a spatial graph convolutional method for predicting protein interfaces. (Yang et al., 2020; Torng & Altman, 2019) use a graph auto-encoder for protein-protein interaction (PPI) classification. (Wang et al., 2022) combine a pretrained AA language model with a GCN representing the structure to predict protein fitness(Yang et al., 2022b).

## 3 Empirical Study

### Tasks and Datasets

We carry out empirical experiments on two function prediction tasks: Enzyme Commission (EC) Numbers prediction (Bairoch, 2000) and Gene Ontology (GO) Terms prediction (Ashburner et al., 2000), following (Gligorijević et al., 2021).

Specifically, we predict levels 3 and 4 of the EC scheme. GO divides proteins into three hierarchically related functional classes, namely Molecular Function (MF), Biological Process (BP), and Cellular Component (CC). We noticed that the GO annotations in SIFTS are incomplete, some proteins have certain functions but are not included in SIFTS, so we process the GO annotations (Dana et al., 2019) of all protein chains in PDB and Swiss-Prot to obtain a binary classification dataset for each single function; the data statistics are shown in Appendix 5.1. In this work, we only evaluate performance on metal ion binding and nucleic acid binding. We use the filtering method from (Gligorijević et al., 2021) to ensure that the sequence similarity between the test set and the training set is no more than 95%. The structures used in this paper are deposited in the PDB before December 21, 2021.

#### 3.1 Graph Construction

In this section, we propose to formulate protein structure modeling by graphs (i.e. graph construction (GC)) using three ways: (1) residue-level GC (RGC) utilizes only the coordinates of the *C_α_* atom of each residue, and represent the coordinate of each residue as nodes; (2) backbone atom-level GC (BAGC) employs 4 atoms (*C_α_*, *C*, *N*, *O*) of each residue and represents them as nodes; (3) all atom-level GC (AAGC) that models all atoms of each residue, including both backbone and side chains, and represents atoms as nodes. The process of protein GC is shown in Figure 1.

##### Definition 3.1

(**Protein Graph**) A protein graph (PG) is defined as 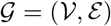, where 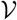 and 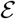 are node set and edge set respectively. Each node is a residue or an atom according to the specific GC method. An edge is constructed when the distance of two nodes are less than a threshold value ε.

**Figure 1:**
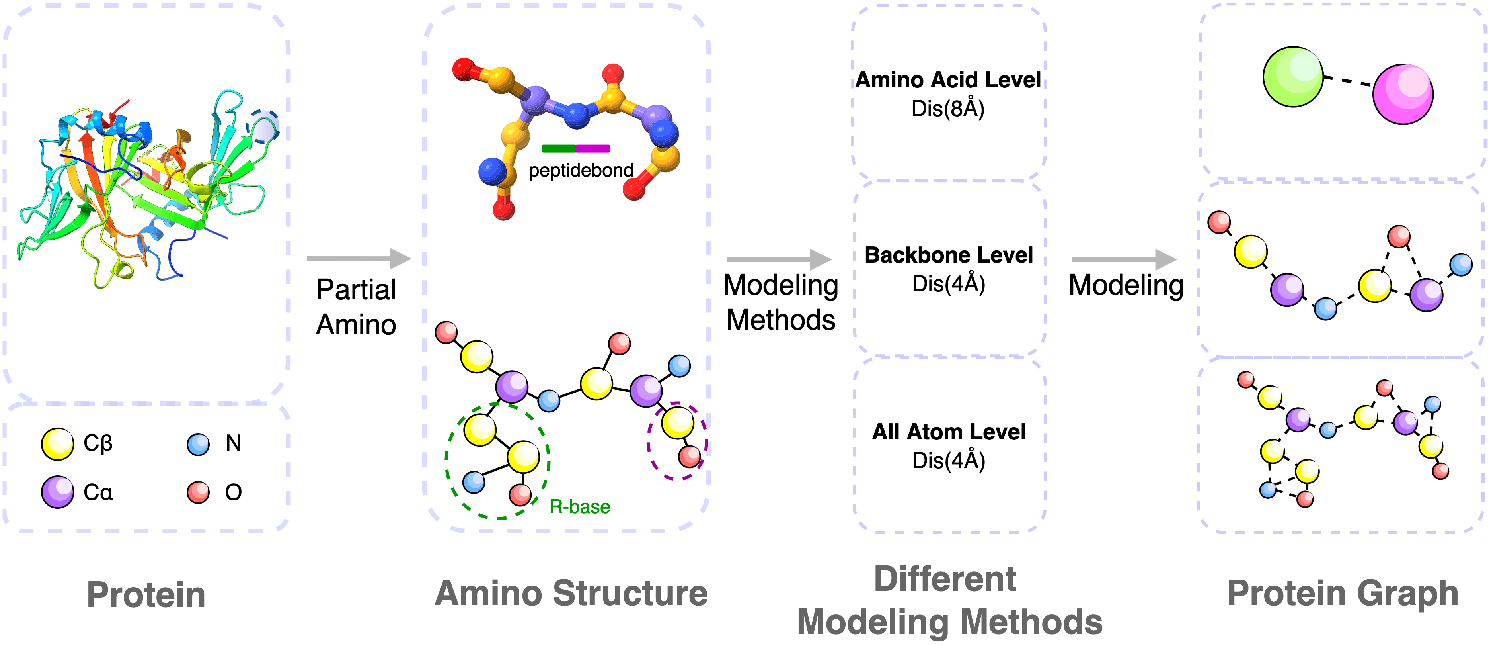
Schematic of modeling protein 3D structures by three types of granularities. The Protein Graph part shows three graphs constructed by amino acids, backbone atoms, and all atoms from top to bottom.

### RGC

For RGC, nodes are residues and edges are the contact information between residues in the 3D space. Here we try two mapping methods, the first RGC_AT is that we define two residues having contact when the distance between the two *C_α_* atoms of them is less than a threshold value *ε*. *ε* is empirically set to 8Å in this paper.^2^.

Each node’s structural input features consist of three dihedral angles of the protein backbone (*ϕ_i_*,*ψ_i_*,*ω_i_*) and embed these on the 3-torus as {sin, cos} × (*ϕ_i_*,*ψ_i_*,*ω_i_*), also we have include the amino acid type as part of the node feature. The input edge features for the *i^th^* residue consist of the dihedral and planar angles and the Euclidean distance between the *C_α_* atom of residue i and the *C_α_* atoms of its distance less than a threshold value 8Å. The construction method of the dihedral angle is shown in the Figure 2.

**Figure 2:**
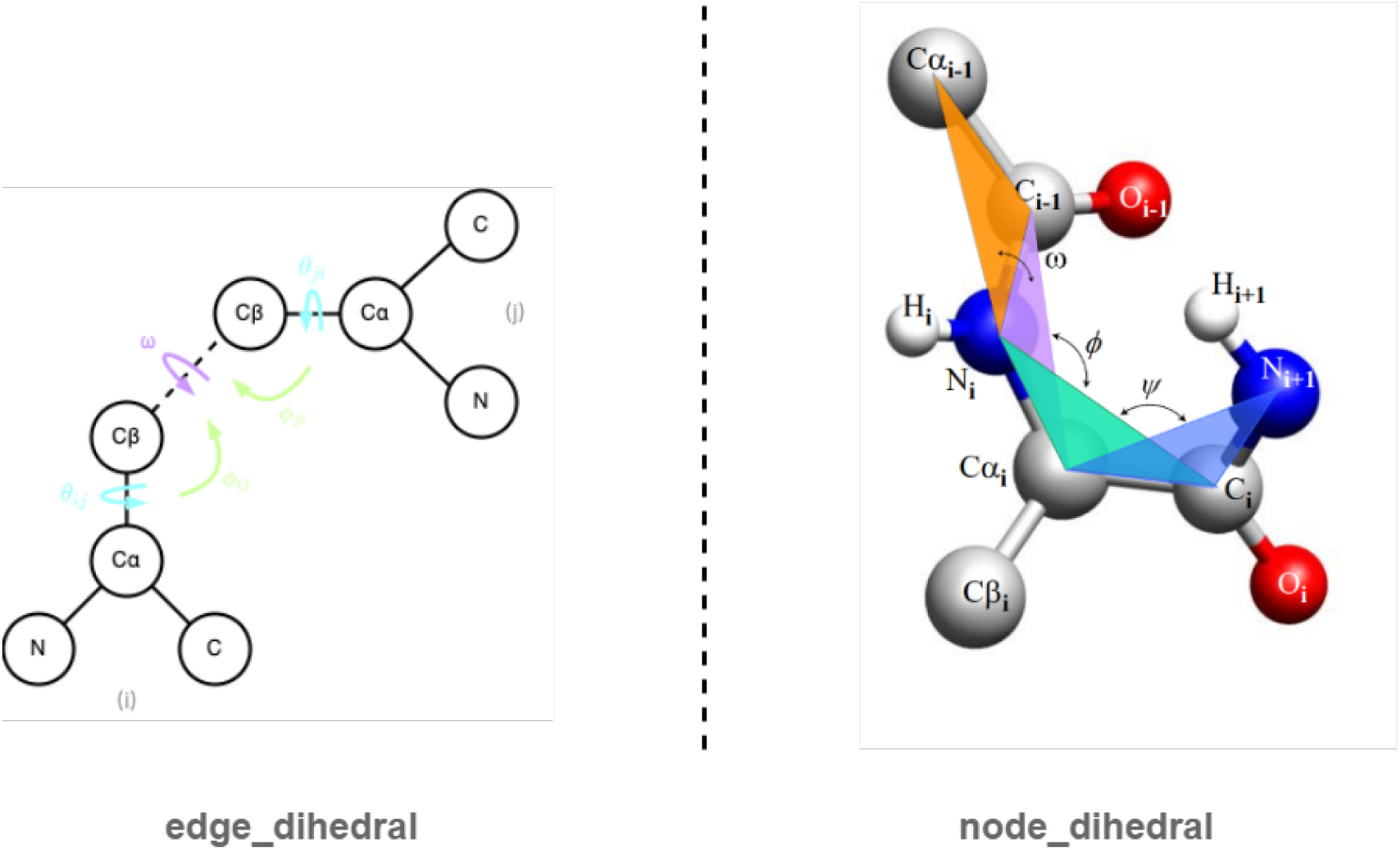
Two different methods for calculating dihedral angles. The left is the calculation of the dihedral angle based on the spatially adjacent amino acids. The right is the calculation of the dihedral angle based on the adjacent amino acids in the sequence.

The second RGC TN method, the node features are the same as in RGC AT. For the edge features, we refer to the distance, direction, and orientation construction methods in (Ingraham et al., 2019) paper.

### BAGC

Intuitively, RGC is coarse-grained relative to the atomic-level GC. Hence, we attempt to explicitly model atoms in the backbone of a residue, which includes *C_α_*, *C*, *N*, *O*. We treat each atom as a graph node. The main difference from RGC is that the distance threshold *ε* that applies to RGC is too large for the atomic-level modeling. It has been experimentally tested that *ε* = 4 Å is better-suited to the atom-level GC.

### AAGC

Instead of using only the backbone atoms, we take account of all atoms in both the main chain and side chain. This is heuristic since the unique property of amino acids is determined by their side chain. Similar to BAGC, each atom is treated as a node, and if *ε* between two nodes is less than 4 Å, an edge will be added between the two nodes.

#### 3.2 Graph Neural Networks (GNN)

GNN is a natural fit to protein 3D structures. Yet it is unknown, among so many top-performed GNN models, which one performs the best when predicting protein various functions. We use 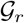, 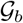, 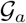 to represent graphs constructed by RGC, BAGC and AAGC respectively. Denote 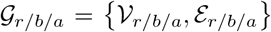, where 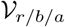, 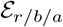 are sets of vertices and edges respectively. Each element in 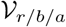 and 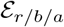 is a vertex and edge generated based on the above three GC methods. We learn protein structure representations through GNNs, and in our model the GNN layer is defined as

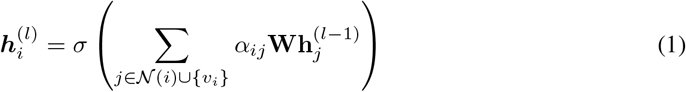

***W*** is a learnable weight matrix, ***a**_ij_* represents the attention correlation coefficient, *j* ∈ *N_i_* represents traversing all nodes adjacent to node i and ***v**_i_* represents node i itself. For more detail on using and not using dihedral angles, distances, etc., the definition of ***a**_ij_* is described in the Appendix A.25.1.

##### Evaluation

We investigated the performance of three common GNN architectures on four datasets related to GO (CC), EC, metal ion binding, and nucleic acid binding. To evaluate the models, We choose two widely adopted metrics (Zhou et al., 2019): maximum protein center F score *F_Max_* and pin-center area under the exact recall curve *AUPR_pair_*, as shown in Table 2. Interestingly, we found that no single model consistently performed best on four tasks. GAT yielded relatively better results overall, except for the F_Max metric in the GO prediction task. This is reasonable, given that these models have shown similar performance in many traditional machine learning tasks.

**Table 1:** Dataset descriptions

| Dataset | Train | Validation | Test |
| --- | --- | --- | --- |
| EC | 16778 | 1865 | 2071 |
| CC | 29902 | 3323 | 3416 |
| Metal ion binding | 72692 | 10000 | 10000 |
| Nucleic acid binding | 52380 | 5000 | 5000 |

**Table 2:**
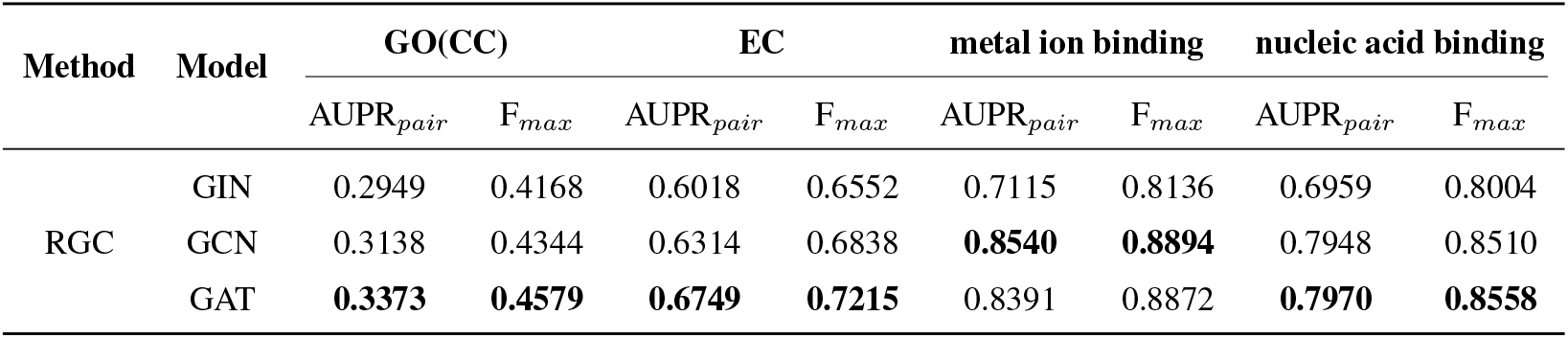
Results of three GNN architectures on four datasets

We conducted a comparison of graph construction methods on the same four datasets presented in Table 3. Overall, we observed that residue-level protein graphs constructed using RGC do not underperform atomic-level graphs constructed using BAGC and AAGC, despite containing less information. Furthermore, the addition of backbone dihedral angles improved performance. The primary difference between the RGC_TN and RG_AT methods is that the former employs a transformer network and incorporates direction, orientation, and distance distribution information in the edge features, while the latter only includes distance and dihedral angle information.

**Table 3:** Results of protein function predictions. ACC is a standard metric for evaluating binary classification.

| Method | Model | GO(CC) |  | EC |  | metal ion binding |  | nucleic acid binding |  |
| --- | --- | --- | --- | --- | --- | --- | --- | --- | --- |
|  |  | AUPR<sub>pair</sub> | F<sub>max</sub> | AUPR<sub>pair</sub> | F<sub>max</sub> | AUPR | ACC | AUPR | ACC |
| BAGC | GAT | 0.3140 | 0.3216 | 0.5699 | 0.5950 | 0.8253 | 0.8618 | 0.7850 | 0.8468 |
| AAGC | GAT | 0.3228 | 0.3285 | 0.5700 | 0.6030 | 0.8416 | 0.8634 | 0.7969 | 0.8506 |
| RGC_AT | GAT | **0.3373** | **0.4579** | 0.6749 | 0.7215 | 0.8391 | 0.8872 | 0.7970 | 0.8558 |
| RGC_TN | GTN | 0.3287 | 0.4506 | **0.8017** | **0.8195** | **0.8593** | **0.8959** | **0.8617** | **0.8972** |

In Appendix 5.3, we present the results of ablation experiments conducted on the RGC_AT and RGC_TN methods. These experiments highlight the importance of domain-specific features and address our **Q1: whether residue-level graph construction with direction, orientation, and distance distribution features are favorable for current leading GNN models. Our findings suggest that such features are indeed beneficial.**

#### 3.3 AlphaFold DB vs PDB

AlphaFold has achieved remarkable success in predicting protein 3D structures with near-atomic accuracy. However, it remains unclear whether the structures generated by AlphaFold are sufficiently accurate for use as input in protein representation models. To address this question, we compared the performance of protein graph models using experimental structures from the PDB with those generated by AlphaFold.

The results presented in Table4 are lower than those shown in Table3. This is because when we screened the 200 million protein database predicted by AlphaFold for protein structures corresponding to CC and EC, we found that some proteins were missing. To ensure consistency between the AlphaFold predictions and the PDB data, we removed the missing proteins from the CC and EC data. Additionally, when we initially began this task, AlphaFold had not yet released the 200 million protein structures. Therefore, we evaluated the metal ion binding task using protein structures

**Table 4:** Result comparison of the structures from PDB and AlphaFold on the CC and EC function prediction task.

| Method | Databases | CC |  | EC |  |
| --- | --- | --- | --- | --- | --- |
|  |  | AUPR<sub>pair</sub> | F<sub>max</sub> | AUPR<sub>pair</sub> | F<sub>max</sub> |
| RGC_TN | Alphafold2 | 0.3012 | 0.4405 | 0.7537 | 0.7856 |
|  | PDB | 0.3115 | 0.4438 | 0.7622 | 0.7956 |

generated by AlphaFold.^3^ Further information on the data volume of this task and the results of the metal ion binding task are provided in Appendix5.4.

Through our comparison of PDB and AlphaFold results on CC and EC function prediction tasks, we can confidently conclude that **the 3D structural data predicted by AlphaFold is on par with the data obtained from PDB in terms of its usefulness as input for a protein function predictor.**

#### 3.4 Pretrain Strategy

Sequence-based pre-training has proven to be highly effective for protein language models, such as ESM-1b and ProtBert. However, it remains unclear whether pre-training directly on protein structures can lead to similar success. Recently, AlphaFold DB has released around 200 million high-precision protein structures. This motivates us to evaluate pre-training effects on these generated protein structures.

##### Experiment Setup

We utilized the popular masked token pre-training method (Devlin et al., 2018) to pre-train protein structural data. During each training step, we randomly mask the attributes of graph nodes with a 15% probability. The model then learns the representation of the masked node through its neighbor nodes and predicts the attributes of the masked node accordingly. As previously mentioned, all datasets used in this experiment were obtained from AlphaFold DB. Initially, we selected proteins with a plddt score greater than 80 from the 200 million proteins in the database. We then employed mmseqs to filter proteins with a sequence similarity greater than 30% and obtained 2 million proteins with corresponding structures for pre-training.

We evaluate the performance of pre-training by zero-shot prediction (ZSP) of the effects of mutations, strictly following (Meier et al., 2021). The reason is because the ZSP task can be directly formulated as the mask token prediction task. We use the Spearman metric (Meier et al., 2021) to evaluate how accurate the predicted tokens are.

##### Results and Discussion

We perform evaluation on BLAT_ECOLX_Ostermeier2014(B2014), BLAT_ECOLX_Ranganathan2015(B2015) and YAP1_HUMAN_Fields-singles(YAP1) datasets. The results are shown in Table 5. We observe that the Spearman correlation is very low, achieving about 0.28 on B2014, 2.23 on B2015, and 0.28 on YAP1. By contrast, it is over 0.7 on B2014 and B2015, and 0.44 on YAP1 by the sequence-based method, i.e. ESM1v (Meier et al., 2021). Although we pre-trained on the CATH 2w(Ingraham et al., 2019) data later, we found that the structure-based method has improved, but it still does not exceed the sequence-based method. In order to intuitively and fairly explore whether the structural information is useful, we removed the distance, direction and orientation structure information in the structure-based method, and found that the Spearman index dropped a lot.

**Table 5:** Results of sequence-based vs structure-based methods on the mutation effect prediction task.

| Datasets | Method | Category | Model | B2014 | B2015 | YAP1 |
| --- | --- | --- | --- | --- | --- | --- |
| AF2 | RGC_AT | Structure | GAT | 0.106 | 0.099 | 0.082 |
|  | RGC_TN | Structure | GTN | 0.286 | 0.234 | 0.280 |
|  | RGC_TN(no_structure) | Sequence | GTN | 0.182 | 0.156 | 0.131 |
| CATH | RGC_TN | Structure | GTN | 0.443 | 0.416 | 0.312 |
|  | RGC_TN(no_structure) | Sequence | GTN | 0.166 | 0.155 | 0.125 |
| UniRef50 | ESM-1V | Sequence | Transformer | 0.706 | 0.707 | 0.441 |

In summary, **current state-of-the-art graph neural network (GNN) models pre-trained on protein structure data are still significantly less effective than sequence-based protein language models (PLMs). However, despite this limitation, structural information remains highly valuable for mutation-related tasks.**

#### 3.5 Sequence vs Structure

In our study, we compare and benchmark sequence-based and structure-based learning methods for the PFP task. For the sequence-based models, we evaluate the two largest and most powerful models in the literature: ESM-1b and ProtBert. For the structure-based models, we assess GAT, GTN, and their pretrained counterparts.

##### Experiment Setup

Currently, there are numerous models for protein sequence representation learning. In our study, we selected ESM-1b, ProtBert, and ProtElect sequence models to compare against structure-based models. In Section 3.1, during the composition method comparison experiment, we discovered that the best choice for representing protein structure is to add upper dihedral angle, distance, direction, and orientation features based on the amino acid level. Consequently, for the sequence and structure comparison experiments, we chose this amino acid-based structural model to compare against the sequence-based model.

To conduct a comprehensive comparison of sequence-based and structure-based protein representation learning, we evaluated models both with and without pretraining. Given that the ESM-1b model based on sequence has significantly more parameters than the GNN model based on structure, we narrowed down the parameters of the original ESM-1b model to match the basic graph model parameters. We then re-pretrained a smaller ESM-1b model.

##### Results and Discussion

According to Table 6, in experiments conducted without pre-training, we observed that the structure-based model outperformed the sequence-based model in both multilabel and single-label binary classification datasets. However, after pre-training, the sequence-based model exhibited a significant improvement in performance and outperformed the structure-based model. In contrast, the structure-based model showed no notable improvement after pre-training. Therefore, we can conclude and answer our **Q3: although protein structure is intuitively more closely related to function than its sequence, the current sequence-based method is still superior to the structure-based method in the PFP task.**

**Table 6:**
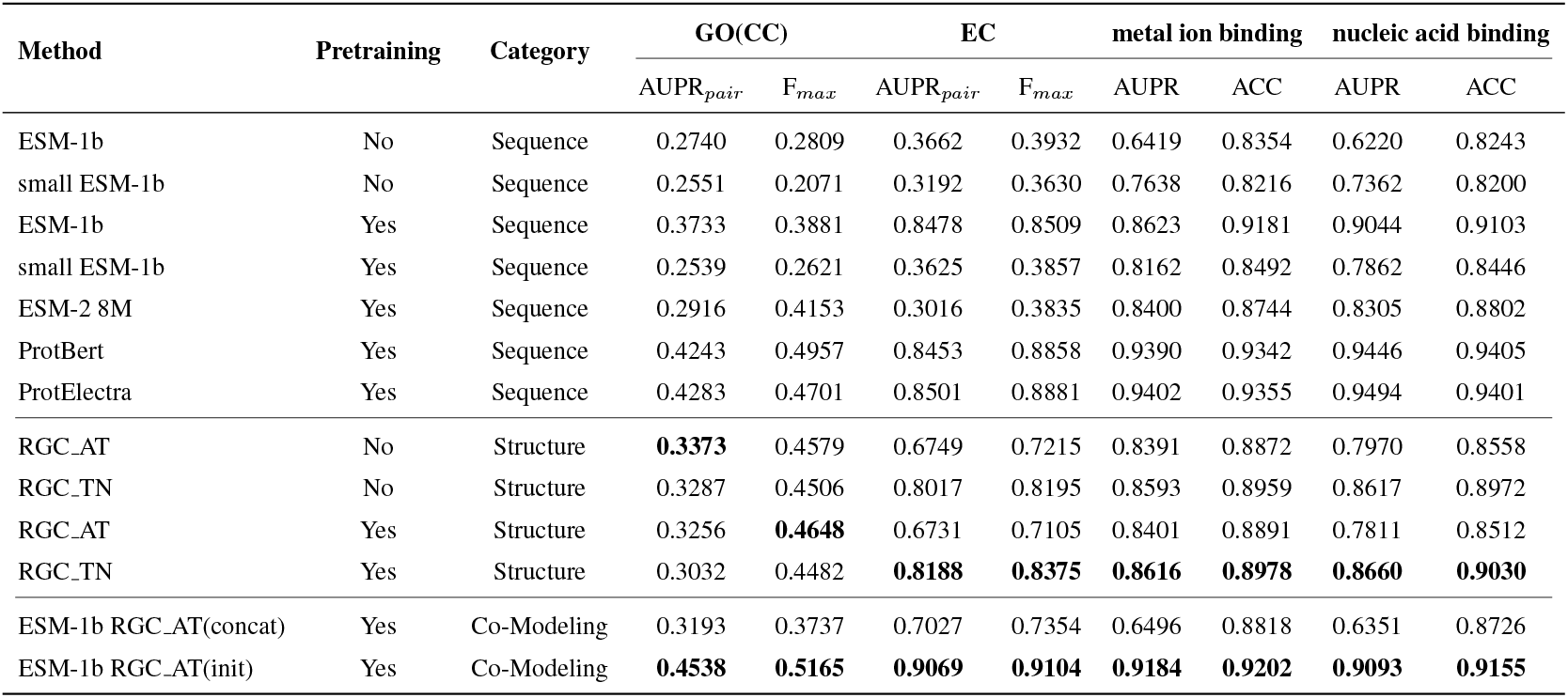
Comparison of different obfuscations in terms of their transformation capabilities

| Method | Pretraining | Category | GO(CC) |  | EC |  | metal ion binding |  | nucleic acid binding |  |
| --- | --- | --- | --- | --- | --- | --- | --- | --- | --- | --- |
|  |  |  | AUPR<sub>pair</sub> | F<sub>max</sub> | AUPR<sub>pair</sub> | F<sub>max</sub> | AUPR | ACC | AUPR | ACC |
| ESM-1b | No | Sequence | 0.2740 | 0.2809 | 0.3662 | 0.3932 | 0.6419 | 0.8354 | 0.6220 | 0.8243 |
| small ESM-1b | No | Sequence | 0.2551 | 0.2071 | 0.3192 | 0.3630 | 0.7638 | 0.8216 | 0.7362 | 0.8200 |
| ESM-1b | Yes | Sequence | 0.3733 | 0.3881 | 0.8478 | 0.8509 | 0.8623 | 0.9181 | 0.9044 | 0.9103 |
| small ESM-1b | Yes | Sequence | 0.2539 | 0.2621 | 0.3625 | 0.3857 | 0.8162 | 0.8492 | 0.7862 | 0.8446 |
| ESM-2 8M | Yes | Sequence | 0.2916 | 0.4153 | 0.3016 | 0.3835 | 0.8400 | 0.8744 | 0.8305 | 0.8802 |
| ProtBert | Yes | Sequence | 0.4243 | 0.4957 | 0.8453 | 0.8858 | 0.9390 | 0.9342 | 0.9446 | 0.9405 |
| ProtElectra | Yes | Sequence | 0.4283 | 0.4701 | 0.8501 | 0.8881 | 0.9402 | 0.9355 | 0.9494 | 0.9401 |
| RGC-AT | No | Structure | **0.3373** | 0.4579 | 0.6749 | 0.7215 | 0.8391 | 0.8872 | 0.7970 | 0.8558 |
| RGC-TN | No | Structure | 0.3287 | 0.4506 | 0.8017 | 0.8195 | 0.8593 | 0.8959 | 0.8617 | 0.8972 |
| RGC-AT | Yes | Structure | 0.3256 | **0.4648** | 0.6731 | 0.7105 | 0.8401 | 0.8891 | 0.7811 | 0.8512 |
| RGC-TN | Yes | Structure | 0.3032 | 0.4482 | **0.8188** | **0.8375** | **0.8616** | **0.8978** | **0.8660** | **0.9030** |
| ESM-1b RGC-AT(concat) | Yes | Co-Modeling | 0.3193 | 0.3737 | 0.7027 | 0.7354 | 0.6496 | 0.8818 | 0.6351 | 0.8726 |
| ESM-1b RGC-AT(init) | Yes | Co-Modeling | **0.4538** | **0.5165** | **0.9069** | **0.9104** | **0.9184** | **0.9202** | **0.9093** | **0.9155** |

Through analysis of the results and the models, we speculate that the structure model did not achieve ideal results because the MLM training method is too simplistic, and the model can only learn the structural representation of the amino acids surrounded by the mask. As a result, its overall protein representation learning ability is not as strong. However, we observed that this training method can be beneficial when performing zero-short mutations.

#### 3.6 Sequence Fusion Structure

Given that 200 million protein structures have been published, significantly increasing the possibility of fusing sequences in large-scale pretrained structures, we speculate that the combination of the two models will outperform sequence-based or structure-based models.

##### Experiment Setup

In our study, we utilized the ESM-1b model based on sequence and the GAT model based on structure. We employed two fusion methods to evaluate their effectiveness. The first method directly splices the output of the ESM-1b model and the GAT model and feeds it to the classifier for the final prediction. The second method involves taking the output of the ESM-1b model as the initialization characteristics of nodes in the graph. The protein characterization obtained is then passed through subsequent GAT and pooling operations before being sent to the classifier for the final prediction. The model structures for both methods can be found in Figure 4.

**Figure 3:**
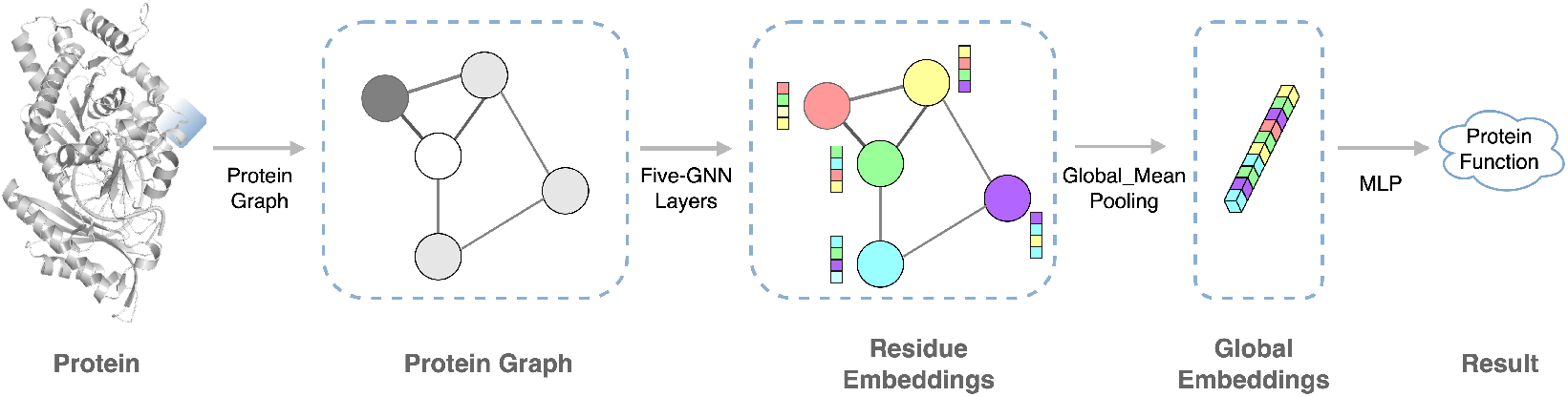
The schematic of graph-based protein function prediction.

**Figure 4:**
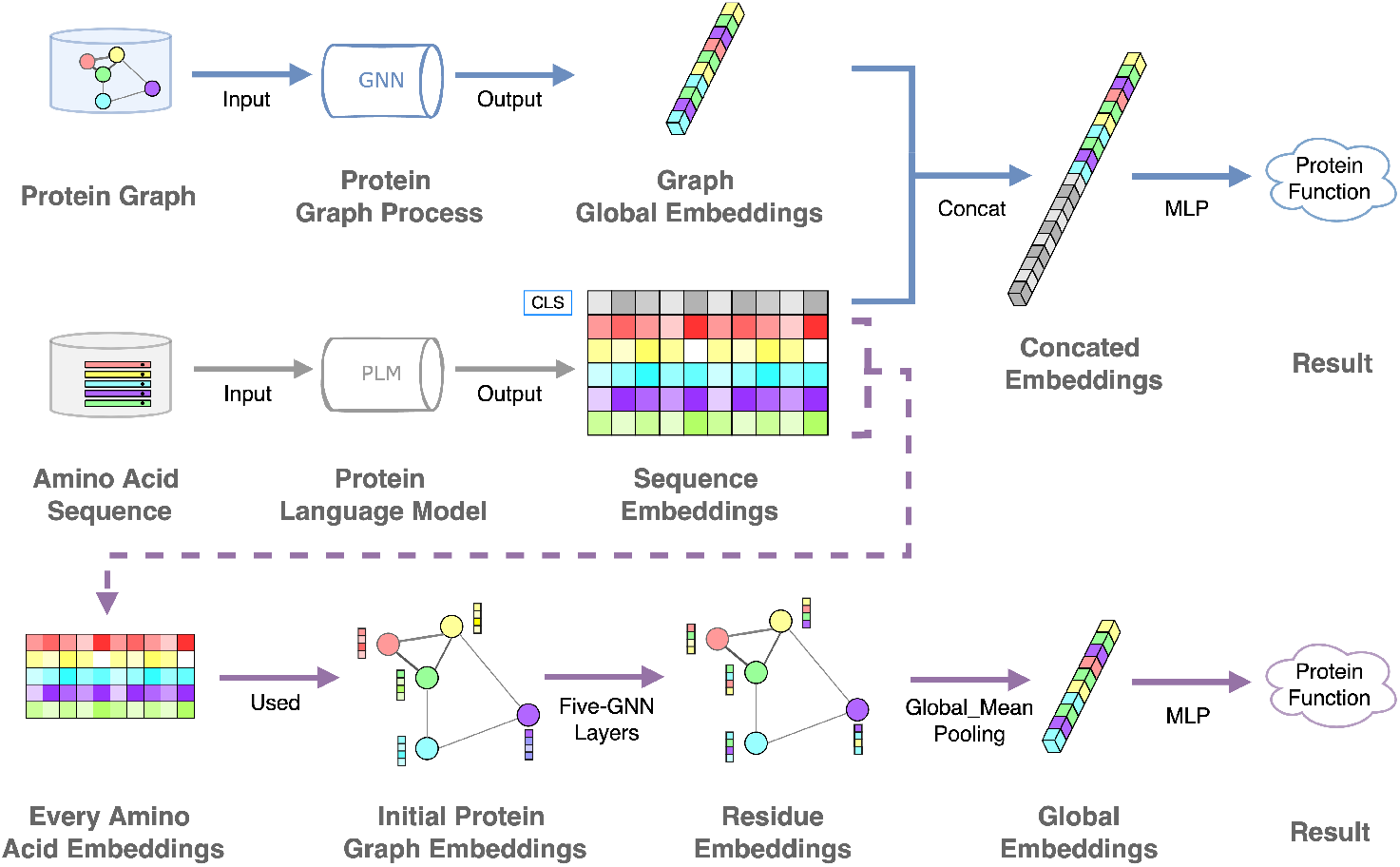
Two methods of sequence-structure fusion methods.

##### Results and Discussion

As shown in Table 6, the fusion method that utilizes ESM-1b as the initial node in the graph achieved the best performance. By integrating sequence and structure, this method outperformed the approach that only leverages either sequence or structure.

## 4 Conclusion

In summary, we initially determined that amino acid hierarchical modeling is the most effective among the three graphical modeling approaches. Next, we evaluated the ability to predict protein function using AlphaFold2-generated protein structures and real protein structures, which confirmed that AlphaFoldDB is a useful tool for predicting protein function. Furthermore, we pre-trained the 200 million protein structures released by AlphaFold2 and compared them with the sequence model. We observed that pre-training significantly improved the effectiveness of the sequence model, while the pre-trained structure model showed no notable impact, which may be due to the simplicity of the MLM training method. Lastly, we discovered that fusion methods yield better results than using only sequences or structures.

**Figure 5:**
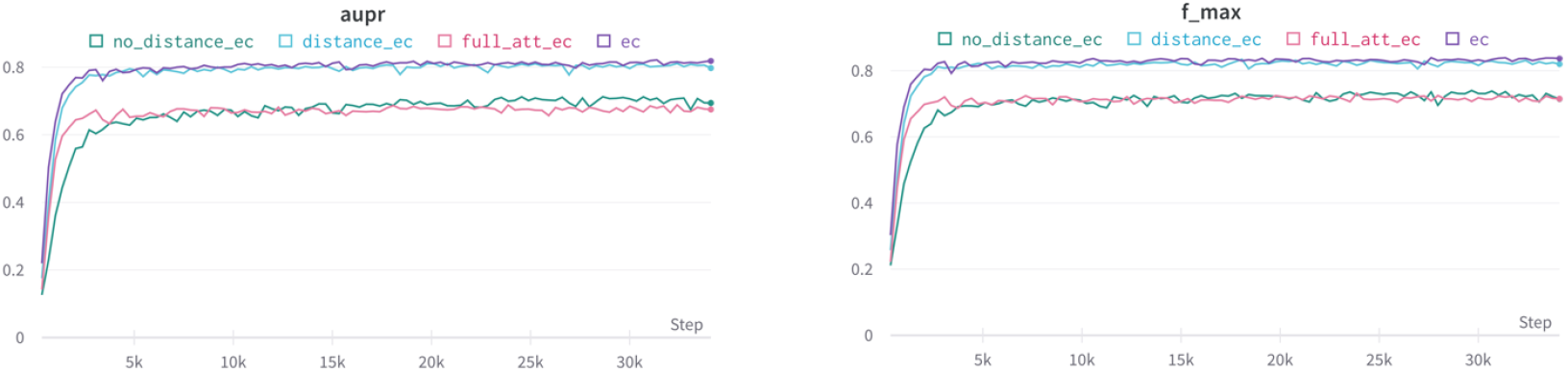
The ablation results of the distance and orientation features on the ec dataset in the RGC_TN method.

**Figure 6:**
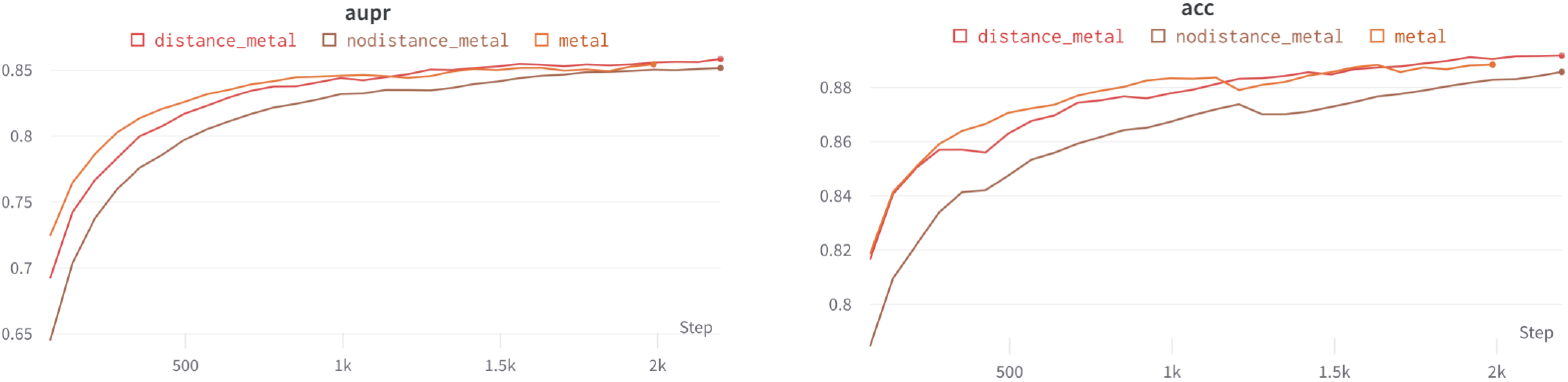
The ablation results of the distance and orientation features on the metal ion binding dataset in the RGC TN method.

**Figure 7:**
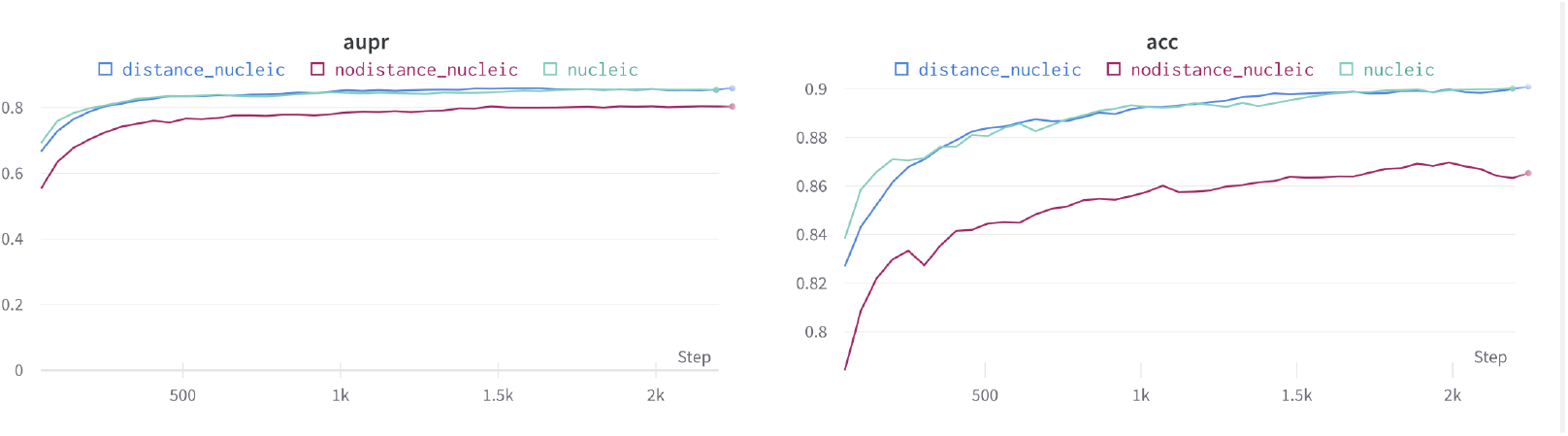
The ablation results of the distance and orientation features on the nucleic acid binding dataset in the RGC TN method.

**Table 7:** Statistics for the number of positive samples for each function in 3 GO categories

| GO Categories | Count | Min | Q1 | Median | Q3 | Max | Mean |
| --- | --- | --- | --- | --- | --- | --- | --- |
| Molecular Function | 5578 | 1 | 12 | 45 | 198 | 370325 | 1099 |
| Biology Process | 2819 | 1 | 23 | 121 | 670 | 145904 | 1668 |
| Cellular Component | 2099 | 1 | 20 | 99 | 710 | 440670 | 3639 |

**Table 8:** The top 10 functions with largest number of positive samples in each GO categories

| GO categories | Function | Count |
| --- | --- | --- |
| **Molecular Function** | binding | 370325 |
|  | catalytic activity | 277662 |
|  | organic cyclic compound binding | 175874 |
|  | heterocyclic compound binding | 174491 |
|  | protein binding | 170131 |
|  | nucleic acid binding | 160253 |
|  | ion acid binding | 149639 |
|  | cation acid binding | 147506 |
|  | metal ion binding | 147334 |
|  | hydrolase activity | 114149 |
| **Biology Process** | cellular process | 145904 |
|  | response to stimulus | 90091 |
|  | biological regulation | 88743 |
|  | regulation of biological process | 73316 |
|  | response to chemical | 70574 |
|  | metabolic process | 68315 |
|  | cellular component organization or biogenesis | 65628 |
|  | viral process | 64613 |
|  | cellular component organization | 60127 |
|  | regulation of cellular process | 52665 |
| **Cellular Component** | cellular anatomical entity | 440670 |
|  | intracellular anatomical structure | 343227 |
|  | organelle | 277881 |
|  | cytoplasm | 272567 |
|  | intracellular organelle | 266881 |
|  | membrane-bounded organelle | 198593 |
|  | intracellular membrane-bounded organelle | 189133 |
|  | membrane | 185371 |
|  | protein-containing complex | 164447 |
|  | cytosol | 143511 |

**Table 9:** AlphaFold2 VS PDB task Dataset description

| Dataset | Train | Validation | Test |
| --- | --- | --- | --- |
| EC | 11649 | 1457 | 1542 |
| CC | 24569 | 2712 | 2936 |
| Metal ion binding | 8000 | 1000 | 1000 |

**Table 10:**
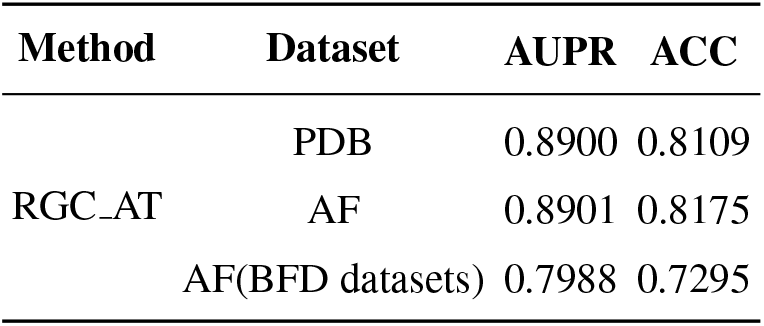
Result of the metal ion binding function prediction task.

**Table 11:**
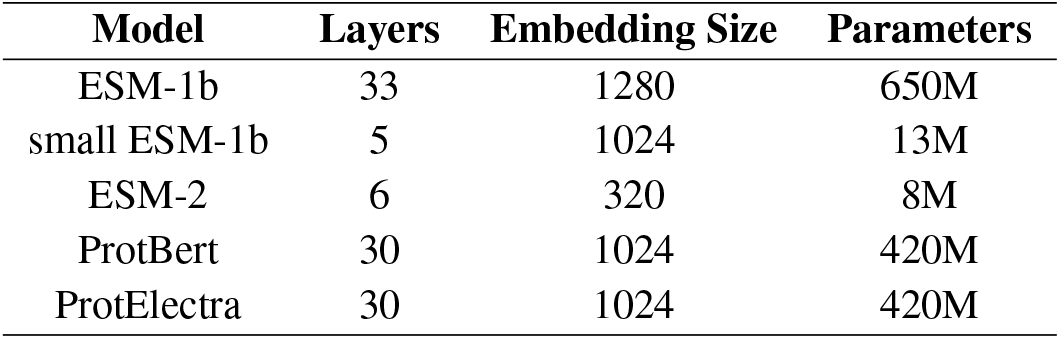
Model Details

| Model | Layers | Embedding Size | Parameters |
| --- | --- | --- | --- |
| ESM-1b | 33 | 1280 | 650M |
| small ESM-1b | 5 | 1024 | 13M |
| ESM-2 | 6 | 320 | 8M |
| ProtBert | 30 | 1024 | 420M |
| ProtElectra | 30 | 1024 | 420M |

<sup>4</sup>https://bfd.mmseqs.com/

**Table 12:** Results of protein function predictions. ACC is a standard metric for evaluating binary classification.

| Method | GO(CC) |  | EC |  | metal ion binding |  | nucleic acid binding |  |
| --- | --- | --- | --- | --- | --- | --- | --- | --- |
|  | AUPR<sub>pair</sub> | F<sub>max</sub> | AUPR<sub>pair</sub> | F<sub>max</sub> | AUPR | ACC | AUPR | ACC |
| RGC_AT | 0.3243 | 0.4172 | 0.5783 | 0.6390 | 0.8319 | 0.8801 | **0.8025** | 0.8543 |
| RGC_AT(edge_dihedral) | 0.3212 | 0.4237 | 0.5914 | 0.6533 | 0.8291 | 0.8748 | 0.7889 | 0.8477 |
| RGC_AT(edge_distance_dihedral) | 0.3237 | 0.4408 | 0.6141 | 0.6752 | 0.8318 | 0.8764 | 0.8011 | 0.8551 |
| RGC_AT(node_dihedral) | 0.3240 | 0.4462 | 0.6538 | 0.7044 | 0.8311 | 0.8795 | 0.7956 | 0.8559 |
| RGC_AT(node_edge_dihedral) | **0.3373** | **0.4579** | **0.6749** | **0.7215** | **0.8391** | **0.8872** | 0.7970 | **0.8558** |

## 5 Appendix

## 5.1 Binary Classification datasets description

We process the GO annotations of all protein chains in PDB and Swiss-Prot to obtain a binary classification datasets for each single function. The statistics of positive samples number of each function in 3 categories are shown in table. A positive sample is a single protein chain who have this certain function.

We show the top 10 functions with the largest number of positive samples in each GO categories in table8.

## 5.2 GNN layer description

If we do not use information such as dihedral angle and distance on the edge, then ***a**_ij_* is defined as

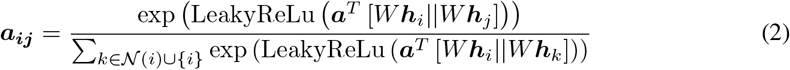

But if we use information such as dihedral angle and distance on the edge, then ***a**_ij_* is defined as

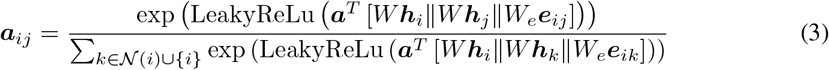

## 5.3 Amino acid mapping ablation experiment

The ablation results of features such as dihedral angle and distance in the RGC_AT method are shown in the table 12.The ablation experiments of the RGC_TN method are shown in Figures 5, 6, and 7. We can find that the distance feature plays a very important role in the classification task. In addition, in the edge relationship in the RGC_TN method, we only select the top 30 nodes with the closest distance. When we removed this restriction and turned into a fully connected graph, we found that the results became worse. This can be seen from Figure 5.

## 5.4 Alphafold2 vs PDB

Table 10 reports the results using data extracted from PDB and generated by AlphaFold. We observe that when searching MSA from only one database (i.e. the BFD^4^ database) GNN with the structural data generated by AlphaFold yields worse results than that using the gold standards from PDB. However, by searching MSA from all four databases, the results are obviously improved and on par with using data from PDB.

## 5.5 Training Detail

The training details of all models appearing in our paper are shown in the Table11 below, we try to keep the training environment of all models the same.

## Footnotes

1 https://www.uniprot.org/help/uniprotkb

2 We have experimented with several other values and found that *ε* = 8Å worked best.

3 It is worth noting that generating protein structures using AlphaFold is still a time-consuming process, as it takes approximately 10-30 minutes to predict a given protein on a 48G GPU (graphics processing unit).

4 https://bfd.mmseqs.com/

